# The adolescent pattern of sleep spindle development revealed by HD-EEG polysomnography

**DOI:** 10.1101/2021.12.20.473516

**Authors:** Gábor Bocskai, Adrián Pótári, Ferenc Gombos, Ilona Kovács

**Author notes:** **Author Contributions:** IK and FG designed the experiments; FG, AP and GB analysed the data; all authors contributed to writing the paper.

## Abstract

Sleep spindles are developmentally relevant cortical oscillatory patterns; however, they have mostly been studied by considering the entire spindle frequency range (11 to 15 Hz) without a distinction between the functionally and topographically different slow and fast spindles, using relatively few electrodes and analysing wide age-ranges. Here, we employ HD-EEG polysomnography in three age-groups between 12 to 20 years of age, with an equal distribution between the two genders, and analyse the adolescent developmental pattern of the four major parameters of slow and fast sleep spindles. Most of our findings corroborate those very few previous studies that also make a distinction between slow and fast spindles in their developmental analysis. We find spindle frequency increasing with age, although spindle density change is not obvious in our study. We confirm the declining tendencies for amplitude and duration, although within narrower, more specific age-windows than previously. Spindle frequency seems to be higher in females in the oldest age-group. Based on the pattern of our findings, we suggest that HD-EEG, specifically targeting slow and fast spindle ranges and relatively narrow age-ranges would advance the understanding of both adolescent development and the functional relevance of sleep spindles in general.

## Introduction

Sleep spindles are cortical oscillatory patterns of NREM sleep in the 11 to 15 Hz range originating in the thalamus (Fernandez and Lüthi, 2020). Their functional role in learning (Berencsi et al., 2017) and the development of cognition (Hahn et al., 2019; Reynolds et al., 2018) makes them important from a developmental perspective. We have recently analyzed the topographical distribution of spindle parameters during the adolescent remodeling of the brain and found evidence for a gradual posteriorization of the anatomical localization of fast sleep spindles in late adolescence, revealing an exceptionally long maturational period in humans in this respect (Gombos et al., 2021).

In order to better understand the human development of sleep spindles, and the exact role they might play in the developing brain, it is important to explore the adolescent developmental pattern of the four major parameters of spindles: density - relevant in protecting sleep from sensory input and in memory (Purcell et al., 2017); duration - thought to reflect the level of thalamic inhibition and/or corticothalamic feedback (Rovo et al., 2014); amplitude, reflecting the overall level of thalamocortical recruitment and globality of neural sources involved in rhythmogenesis (Andrillon et al., 2011; Fernandez and Lüthi, 2020); and frequency, reflecting the duration of thalamocortical hyperpolarization-rebound sequences potentially based on myelination (Zhang et al., 2021). Although several papers address age-related patterns in spindle parameter development, some of these do not analyze slow and fast spindles independently, looking at the entire 11 to 15 Hz range e.g., (Kurth et al., 2010; Purcell et al., 2017; Tarokh et al., 2011), or they collapse a wide age-range between childhood and adulthood without specifically analyzing adolescence (Campbell and Feinberg, 2016; Kurth et al., 2010). There are only a handful of studies that make the distinction between slow and fast spindle parameters, and directly analyze those in the adolescent age-range (Goldstone et al., 2019; Hahn et al., 2019; Nader and Smith, 2015; Zhang et al., 2021). However, the shortcoming of these studies is that the number of electrodes is very low (4 to 18), not allowing for topographical precision. Since we have previously shown (Gombos et al., 2021) that there is a bimodal topographical shift of the two spindles in adolescence, this might be a relevant methodological issue since the detection of spindles might depend on electrode placement.

Here we explore the adolescent developmental pattern of the four major parameters of slow and fast sleep spindles with HD-EEG polysomnography employing an individually adjusted method (IAM, Bódizs et al., 2009) of spindle detection in three age-groups between 12 to 20 years of age, with an equal distribution between the two genders. Most of our findings are confirmatory with respect to the pre-existing studies employing only a few electrodes and wide age-groups, and we also find a clear difference between the development of slow and fast spindles in terms of duration, amplitude, and frequency.

## Methods

Our analyses were performed in an existing database of full night polysomnography recordings of 60 young adult and adolescent subjects (Gombos et al., 2021). In that database, subjects were recruited through social media in three age groups, 10 males and 10 females in each age group: 12-year-olds (n=20, mean age=12.45 ± SD .57 years), 16-year-olds (n=20, mean age=15.91 ± SD .48 years), and 20-year-old young adults (n=20, mean age=21.29 ± SD .51 years). Only healthy individuals who were free of sleep and neurological conditions were included in the database. Each subject received a HUF 20,000 (cca. USD 70.00) voucher for participation. The study was approved by the Ethical Committee of the Pázmány Péter Catholic University for Psychological Experiments, and participants/parents gave informed written consent.

Subjects spent two consecutive nights in the sleep laboratory of the Pázmány Péter Catholic University Budapest where 128 channel HD-EEG polysomnography (PSG) was recorded. Participants were asked to maintain their usual sleep-wake cycles for 5 days prior to the study; cooperation was not monitored. Participants were asked not to take any drugs or caffeine on the day of the study and to refrain from taking naps on study days. Subjects went to bed between 10 p. m. and 11.30 p.m. in the sleep laboratory, where they could sleep until they woke up spontaneously in the morning, during this time all night 128 channel HD-EEG, electrooculogram (EOG), and electromyogram (EMG) were recorded. For the procedure of EEG recordings, hardware environment, data processing and sleep spindle extraction method see the study by Gombos et al (Gombos et al., 2021).

This study is based on the analysis of EEG recordings from the second night being free from the effects of the first, adaptation night. The sleep states of 20-second epochs of the whole-night NREM EEG recordings were scored according to standardized criteria (Gombos et al., 2021). Individual slow and fast spindles were detected using the IAM method (Bódizs et al., 2009). The IAM is based on the average amplitude spectrum of NREM sleep. The frequency criterion for slow and fast sleep spindles was derived from the inflection points of individual spindle peaks between 9 and 16 Hz in the spectra. The IAM method was used to determine the individual and lineage-specific density (spindles × min-1), duration (s) and amplitude (µV) of slow, frontally dominant, and fast, centro-parietally dominant sleep spindles. Peak frequencies were determined using the individualized spectral peak frequency centroid (Center of Gravity, CoG) in the spindle range (11-16 Hz). The CoG values were obtained using the method described in a study by Dustman (Dustman et al., 1999). Outliers were excluded from the original data set using Tukey’s fence method (Gombos et al., 2021).

Slow spindles which are frontally dominant were analyzed on averaged measurements of slow spindle values of the 37 channels in the frontopolar and frontal regions (FPF). Fast spindles which are centro-parietally dominant were analyzed on averaged measurements of fast spindle values of the 42 channels in the centro-parietal regions (CP).

We separately analyzed the effects of age and sex on the frequency, amplitude, duration, and density of slow and fast sleep spindles. Sleep spindle amplitude, duration, density and frequency were analyzed as dependent variables in four different GLM (one for each spindle feature) with between subjects’ factors being age group and sex and the within-subject factor being spindle type (slow vs fast). Where we found significant main effects or interaction effects, we applied Fisher LSD tests for post-hoc analyses. If the main effect was not significant, we used the more stringent Tukey post-hoc test, which does not require a significant main effect, to test for differences between subgroups.

For sleep data processing (classification of sleep states, rejection of artifacts and spindle detection using IAM), we used the program FerciosEEGPlus 1.3 (Gombos et al., 2021). For the calculation of CoG values, we used the Microsoft Excel program. All statistical analyses were performed using TIBCO STATISTICA 13.5.0.17 software (TIBCO Software Inc., 2018).

## Results

We broke down the period of adolescence into three age groups 12, 16 and 20 years, and analyzed the full-night EEG data. Sleep spindle amplitude, duration, density, and frequency were analyzed as dependent variables in four different GLM (one for each spindle feature) with between subjects’ factors being age group and sex and the within-subject factor being spindle type (slow vs fast, Figure 1.)

**Figure 1.**
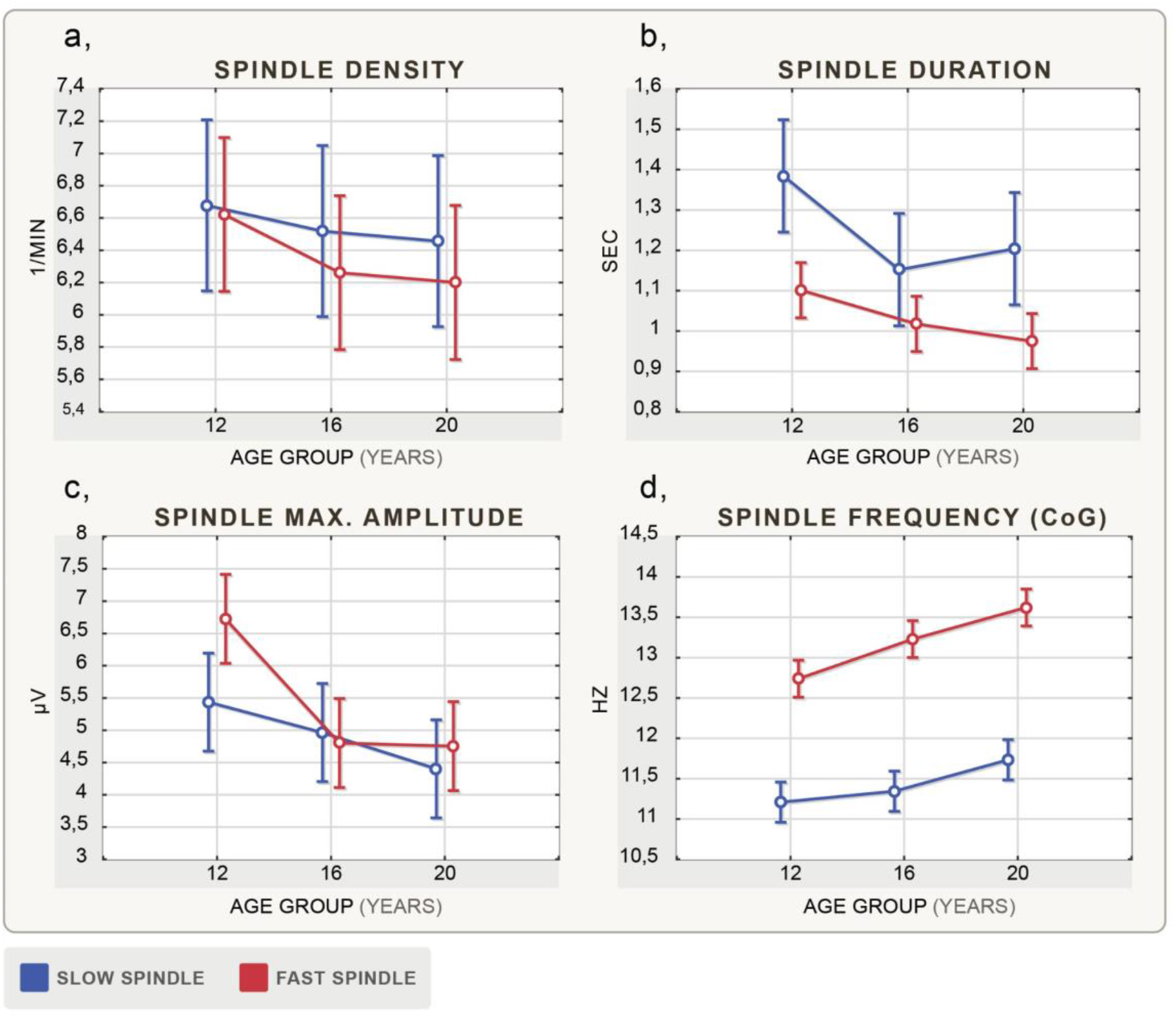
Age-related changes of NREM sleep spindle parameters. Categorical factors: age groups (12 years, 16 years, 20 years), dependent variables: **a**, Spindle density (1/min), **b**, Spindle duration (sec), **c**, Spindle maximum amplitude (μV), **d**, Spindle frequency (Hz). Vertical bars denote 0.95 confidence intervals; the central points of bars represent average values for each age group. Blue color denotes slow spindle, red color denotes fast spindle.

**Figure 2.**
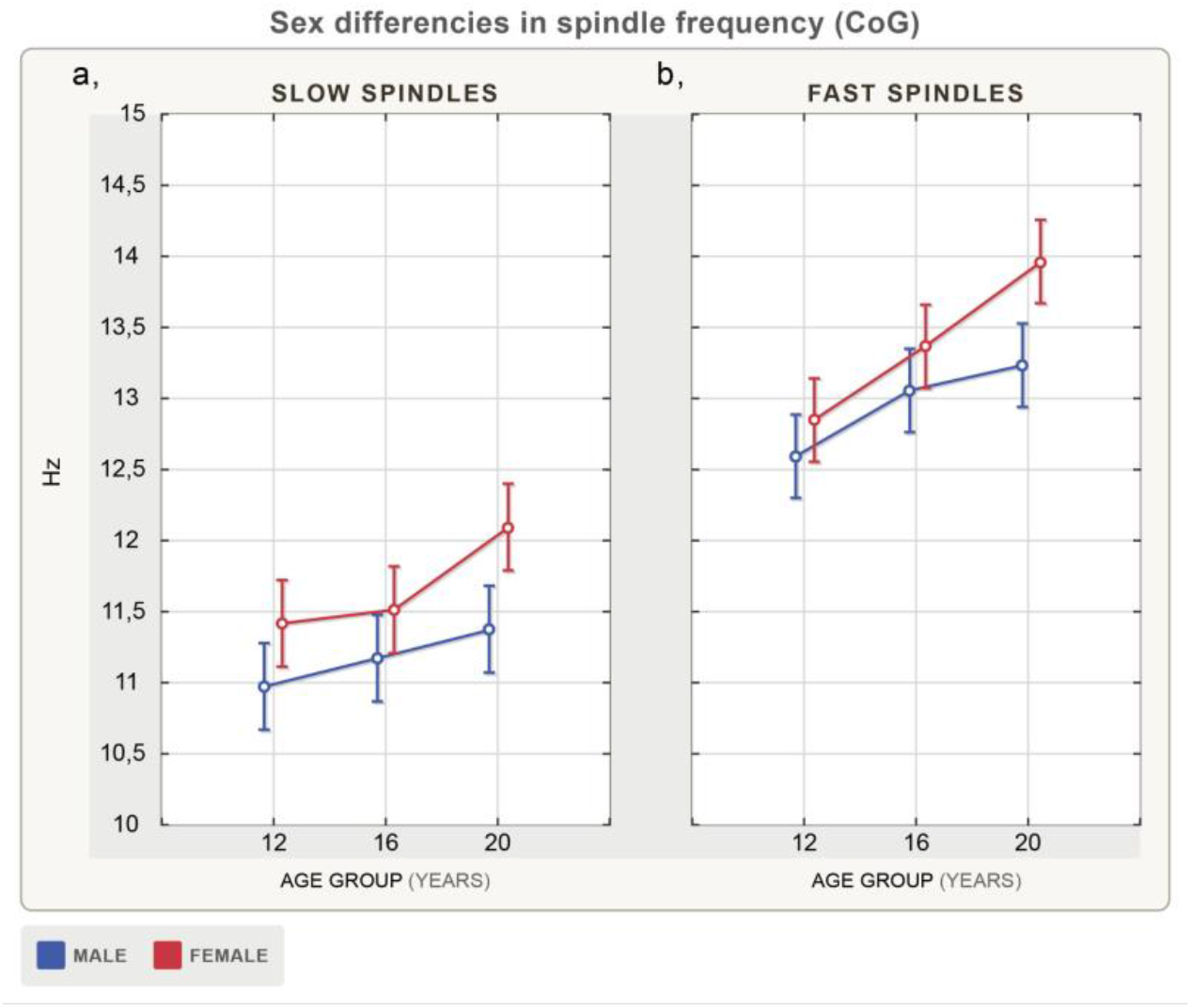
**S**ex and age-related changes (12 years, 16 years, 20 years) of NREM sleep spindle frequency (Hz) is depicted on the left side of the figure for slow spindles, and on the right side for fast spindles. Vertical bars denote 0.95 confidence intervals; the central points of bars represent average values for each age group. Color codes blue: male, red: female.

### Main effects and interactions

The analysis revealed significant main effect of spindle type and age group in terms of duration (F=31.3; df=1; p<.001 and F=4.383; df=2; p=.017, respectively), maximum amplitude (F=5.29; df=1; p=.025 and F=6.422; df=2; p=.003, respectively) and frequency (F=986.32; df=1; p<.001 and F=13.76; df=2; p<.001, respectively). Sex effect was significant only in terms of frequency (F=18.11; df=1; p<.001). The interaction effect of spindle type and age group was significant in maximum amplitude (F=3.937; df=2; p=.025) and frequency (F=4.04; df=2; p=.023). No significant interaction effect of spindle type, sex and age group was found for any of the sleep spindle features.

Where we found significant main effects or interaction effects, we applied Fisher LSD tests for post-hoc analyses. If the main effect was not significant, we used the more stringent Tukey post-hoc test.

### Density

Post-hoc tests revealed no significant differences (Figure 1/a).

### Duration

Post-hoc tests revealed significantly longer sleep spindle duration at age 12 than at ages 16 and 20 (Fisher LSD, p=.012 and p=.015, respectively). Slow spindle duration decreased significantly from age 12 to age 16 (Tukey, p=.037, Figure 1/b).

### Maximum amplitude

We found a significant decrease in terms of maximum amplitude from age 12 to age 16 and from age 12 to age 20 (Fisher LSD, p=.009 and p=.001, respectively). Slow spindle maximum amplitude decreased significantly from age 12 to 20 (Fisher LSD, p=.048). Fast spindle maximum amplitude decreased significantly from age 12 to 16 and 12 to 20 (Fisher LSD, p<.001 for both, Figure 1/c).

### Frequency

Frequency increased with age across the whole age range. Significantly higher spindle frequency was found in 16-year-olds than in 12-year-olds and in 20-year-olds than in 12- and 16-year-olds (Fisher LSD, p=.022, p<.001 and p=.006, respectively). Slow spindle frequency increased significantly from age 12 to age 20, and from age 16 to age 20 (Fisher LSD, p<.001 and p=.011, respectively). Fast spindle frequency was significantly higher in 16-year-olds than in 12-year-olds and in 20-year-olds than in 12- and 16-year-olds (Fisher LSD, p=.002, p<.001 and p=.012, respectively). Females in the 20-year age group have significantly higher slow and fast spindle frequency than males (Tukey, p=.049 and p<.041, Figure 1/d).

## Discussion

We explored the adolescent developmental pattern of the four major parameters of slow and fast sleep spindles with HD-EEG polysomnography in three age-groups between 12 to 20 years of age. Most of our findings corroborate those very few previous studies that also make a distinction between slow and fast spindles in their developmental analysis. With respect to slow spindle density, we were not able to confirm the decline (Nader and Smith, 2015; Zhang et al., 2021), and we only confirm the frequency increase (Zhang et al., 2021) between 16 to 20 years of age. Since the Zhang et al. (2021) study did not specifically target a finer age-resolution in adolescence, nor a very clear distinction between slow and fast spindles, we believe that our data might be more precise. In the case of fast spindles, we confirm the increasing frequency (Goldstone et al., 2019; Zhang et al., 2021), however, we could not confirm the density increase observed by others (Goldstone et al., 2019; Nader and Smith, 2015). With respect to amplitude and duration, we replicate the declining developmental trend of slow spindles (Goldstone et al., 2019; Nader and Smith, 2015; Zhang et al., 2021). The narrower age-window, just as above, might indicate a more precise result.

With respect to sex and age-related changes, we confirm that young adult females tend to have higher spindle frequencies in the case of both slow and fast spindles (Zhang et al., 2021).

In this study, we employed HD-EEG sleep recordings for the first time to investigate the adolescent development of the relevant parameters of both slow and fast sleep spindles. In addition to providing more precision, HD-EEG also provides for the possibility of detailed topographic analysis that seems to be especially relevant in evaluating adolescent sleep recordings. (Gombos et al., 2021). The results confirm some of the earlier findings obtained with fewer electrodes and wider age-windows than in the present study. We also find a clear difference between the development of slow and fast spindles in terms of duration, amplitude, and frequency. Based on the pattern of our findings, we suggest that HD-EEG, specifically targeting slow and fast spindle ranges and relatively narrow age-ranges would advance the understanding of both adolescent development and the functional relevance of sleep spindles in general.

## Acknowledgments

The authors thank T. Jáger for generating the figures and tables. We also thank the adolescents and young adults who participated in this project, and Zoltán György who assisted at all the sleep recordings. This research was supported by the Hungarian National Research, Development and Innovation Office grants NK-104481 and K-134370 to I.K.

## Notes

### Competing Interest Statement

The authors have declared no competing interest.

### Summary of Updates

There was a spelling error in the title. One of the authors changed his mind with respect to authorship. We removed this author from the list of authors.

